# DMC1 attenuates RAD51-mediated recombination in Arabidopsis

**DOI:** 10.1101/2022.07.05.498790

**Authors:** Olivier Da Ines, Jeanne Bazile, Maria E. Gallego, Charles I. White

## Abstract

Ensuring balanced distribution of chromosomes in gametes, meiotic recombination is essential for fertility in most sexually reproducing organisms. The repair of the programmed DNA double strand breaks that initiate meiotic recombination requires two DNA strand-exchange proteins, RAD51 and DMC1, to search for and invade an intact DNA molecule on the homologous chromosome. DMC1 is meiosis-specific, while RAD51 is essential for both mitotic and meiotic homologous recombination. DMC1 is the main catalytically active strand-exchange protein during meiosis, while this activity of RAD51 is downregulated. RAD51 is however an essential cofactor in meiosis, supporting the function of DMC1. This work presents a study of the mechanism(s) involved in this and our results point to DMC1 being, at least, a major actor in the meiotic suppression of the RAD51 strand-exchange activity in plants. Ectopic expression of DMC1 in somatic cells renders plants hypersensitive to DNA damage and specifically impairs RAD51-dependent homologous recombination. DNA damage-induced RAD51 focus formation in somatic cells is not however suppressed by ectopic expression of DMC1. Interestingly, DMC1 also forms damage-induced foci in these cells and we further show that the ability of DMC1 to prevent RAD51-mediated recombination is associated with local assembly of DMC1 at DNA breaks. In support of our hypothesis, expression of a dominant negative DMC1 protein in meiosis impairs RAD51-mediated DSB repair. We propose that DMC1 acts to prevent RAD51-mediated recombination in Arabidopsis and that this down-regulation requires local assembly of DMC1 nucleofilaments.

**Author Summary:** Essential for fertility and responsible for a major part of genetic variation in sexually reproducing species, meiotic recombination establishes the physical linkages between homologous chromosomes which ensure their balanced segregation in the production of gametes. These linkages, or chiasmata, result from DNA strand exchange catalyzed by the RAD51 and DMC1 recombinases and their numbers and distribution are tightly regulated. Essential for maintaining chromosomal integrity in mitotic cells, the strand-exchange activity of RAD51 is downregulated in meiosis, where it plays a supporting role to the activity of DMC1. Notwithstanding considerable attention from the genetics community, precisely why this is done and the mechanisms involved are far from being fully understood. We show here in the plant Arabidopsis that DMC1 can downregulate RAD51 strand-exchange activity and propose that this may be a general mechanism for suppression of RAD51-mediated recombination in meiosis.

## Introduction

To maintain stable ploidy across generations, sexually reproducing eukaryotes must halve the number of chromosomes in gametes with respect to their mother cells. This is accomplished by meiosis, a specialized cellular division in which DNA replication is followed by two sequential rounds of chromosome segregation [1, 2]. Proper chromosome segregation in mitosis, in which a single division follows DNA replication, is ensured by sister-chromatid cohesion established during S-Phase. This however only works once and an additional mechanism is needed in meiosis. Balanced segregation of chromosomes at the first meiotic division relies upon the establishment of physical connections between the two homologous chromosomes. In most species, these connections are realized by reciprocal recombination events called cross-overs (COs). This process also produces novel combinations of alleles from the two chromosomes, recombining the parental genetic contributions in the production of the new generation. Meiotic recombination is thus both essential for fertility and to reshuffle genetic information, with important consequences for genome evolution [1, 2].

Meiotic recombination is initiated by the programmed induction of DNA double strand breaks (DSB) by the SPO11 DNA topoisomerase VI-like complex [3–5] in the chromosomes and their repair by homologous recombination [6–11]. The pair of DNA ends formed are resected by nucleases [12], which remove SPO11 and generate long 3□ single-stranded DNA overhangs (ssDNA), which are bound by the heterotrimeric protein RPA to protect the ssDNA and prevent formation of secondary structures [13–15]. The specialized RAD51 and DMC1 recombinases are then loaded onto the ssDNA to form right-handed helical nucleoprotein filaments [16, 17]. Earlier studies have proposed that recombinase loading occurs asymmetrically, with RAD51 and DMC1 making separate homofilaments on opposite ends of a break [18–20]. However, more recent work in budding yeast and mouse suggested that RAD51 and DMC1 homofilaments do load on both DSB ends resulting in mixed recombinase filaments [21–23]. This mixed recombinase filament organization has been reconstituted *in vitro* [24] and has thus become the favored model for recombinase loading [16, 17, 23–27]. The RAD51/DMC1-DNA nucleoprotein filament is the active molecular species in searching for and invading a homologous DNA template by the 3’-ended DNA strand(s) [16, 17, 26, 27]. This process is the core of the meiotic recombination pathway and results in the formation of a joint recombination intermediate, which can be resolved to yield non-crossover (NCO) or crossover (CO) products [1, 2, 28]. CO between non-sister-chromatids, in combination with sister chromatid cohesion, establish the physical linkages (chiasmata) necessary to ensure proper chromosomal segregation at the first meiotic anaphase [1, 2, 28].

Only CO between non-sister chromatids physically link homologous chromosome pairs and the choice of the sister or non-sister chromatid template for repair of meiotic DSB is thus a key determinant for the outcome of recombination. This must be tightly regulated to favor interhomolog recombination for coordinated chromosomal disjunction at the first meiotic anaphase [29]. With a few exceptions, this crucial step of meiotic recombination requires the co-operation of the two strand exchange proteins, RAD51 and DMC1 (reviewed by [16, 17, 30]). RAD51 and DMC1 emerged following a gene duplication event that occurred in the common ancestor of all eukaryotes [31]. They retain more than 50% amino acid identity and closely related structural and biochemical properties [16, 26, 32–34]. Both enzymes assemble on ssDNA at sites of breaks and promote break repair by searching for and invading an intact homologous DNA template molecule. However, while RAD51 is essential for both mitotic and meiotic homologous recombination, DMC1 is meiosis-specific [16, 27, 35, 36]. Why meiotic recombination requires two strand-exchange proteins is not well understood, but given the similar activities of the two proteins it has been generally assumed that they play complementary roles in catalyzing meiotic DSB repair - with DMC1 promoting interhomolog recombination (IH) and crossing-over, while RAD51 favors sister-chromatid repair of the DSB [18, 37]. This assumption has however been called into question by the characterization of the yeast *rad51-II3A* [38] and Arabidopsis RAD51-GFP proteins [39]. While they retain the ability to form nucleoprotein filaments at DSB, these proteins are unable to catalyze invasion of the homologous template. They do however fully complement the absence of RAD51 in meiosis [38–41]. It is thus the presence of the RAD51 protein and not its strand-exchange activity that is needed in meiosis.

DMC1 is the predominant strand-exchange protein during meiosis, catalyzing most strand-exchange events, while RAD51 acts only to support DMC1 and to repair residual DSBs after IH recombination and synapsis are complete [38, 39, 42–44]. Accordingly, data from budding yeast indicate that RAD51 activity is downregulated during meiosis so that DMC1 catalyzes DNA strand-exchange using the homologous chromosome as a template [38, 45–47]. In budding yeast, down-regulation of Rad51 activity is mediated through inhibition of Rad51-Rad54 complex formation by the coordinated phosphorylation of Hed1 and the Rad51 accessory factor Rad54 by the meiosis-specific kinase Mek1 [18, 45, 46, 48–51]. Hed1 is a meiosis-specific factor that binds to Rad51, blocking access of Rad54 and thereby restricting activity of the Rad51 nucleofilaments in meiosis [48, 49, 51, 52]. Second, phosphorylation of threonine 132 of Rad54 by Mek1 reduces the affinity of Rad54 for Rad51 [47, 50]. Both mechanisms affect Rad51-Rad54 complex formation and this in turn attenuates the activity of Rad51, presumably to minimize the use of sister chromatid templates and hence favor DMC1-dependent inter-homolog recombination.

Apart from budding yeast, the mechanisms suppressing RAD51 activity during meiosis have not been determined. In plants, bioinformatics searches fail to identify homologs of Hed1 and Mek1. Arabidopsis *rad51* mutants are sterile and *dmc1* mutants have severely reduced fertility. They both show defects in pairing and synapsis, but *rad51* and *rad51 dmc1* mutants exhibit extensive SPO11-dependent chromosome fragmentation in meiotic prophase I, while *dmc1* mutants are characterized by the presence of intact univalent chromosomes in meiotic metaphase I [53–55]. Thus, in the absence of DMC1, meiotic DSB are repaired in a RAD51-dependent manner [53, 55–57] which does not generate chiasmata. This has led to the hypothesis that DMC1 could be involved in down-regulation of RAD51 in Arabidopsis [57, 58].

Here, we sought further insights into the mechanism by which RAD51 is downregulated during meiosis in plants and our results support the hypothesis that DMC1 ssDNA binding attenuates RAD51 strand invasion activity. Ectopic expression of DMC1 in vegetative cells blocks RAD51-dependent homologous recombination and renders plants hypersensitive to DNA damage. DNA damage-induced RAD51 focus formation in vegetative cells is not however impaired by ectopic expression of DMC1 and DMC1 also forms damage-induced foci in these cells. The blocking of RAD51 function in vegetative cells thus occurs through inhibition of its activity rather than via inhibition of nucleofilament formation, reminiscent of the situation in meiosis. This conclusion is further supported by the observation that expression of a dominant negative DMC1-GFP fusion protein in meiosis impairs RAD51-mediated meiotic DSB repair. Interestingly, expression of a dominant negative, non-nucleofilament-forming, DMC1-II3A variant did not prevent RAD51-mediated recombination in meiosis or in mitosis, concordant with the argument that the ability of DMC1 to prevent RAD51-mediated recombination requires assembly of DMC1 nucleofilaments at DSB sites. Our data thus suggest that DMC1 triggers down-regulation of RAD51 strand-exchange activity and that local assembly of DMC1 nucleofilaments - but not this strand-exchange activity – may be sufficient to mediate this down-regulation in Arabidopsis.

## Results

### Ectopic expression of DMC1 in vegetative cells

We expressed *DMC1* in vegetative cells, in which RAD51 is normally the only active recombinase, to test the intrinsic ability of DMC1 to downregulate RAD51. The Arabidopsis *DMC1* genomic sequence (*DMC1g*) was cloned and placed under the control of the *RAD51* promoter to drive its expression in somatic cells (Figure 1A). The complementation of the meiotic defects of *dmc1* mutants by this transgene confirms correct expression of functional DMC1 protein (Figure 1B and C). We selected 3 independent *dmc1−/− pRAD51::DMC1g* transgenic lines and show that all were fertile with a number of seeds per silique very similar to that of wild-type plants (Figure 1C and S1 Data). Chromosome spreads of pollen mother cells confirm that the *pRAD51::DMC1g* transgene restores normal meiosis in *dmc1* mutants (S1 Fig). We note that a similar experiment using the *DMC1* coding sequence (*DMC1_CDS_*) instead of the genomic sequence, did not rescue the fertility of *dmc1* mutants, presumably indicating the importance of an intron sequence and/or correct splicing for proper *DMC1* expression (Figure 1B). Knowing that the chimeric *pRAD51::DMC1g* transgene allows high expression of *DMC1* we then tested whether it could drive expression of *DMC1* in somatic cells and whether or not it showed DNA damage-inducible expression, as does *RAD51.* We generated wild-type plants homozygous for the transgene insertion and performed RT-PCR on RNA from one-week-old seedlings. *RAD51* expression is relatively low in wild-type seedlings grown under standard conditions but its expression is strongly induced in seedlings treated with 20 µM Mitomycin C (MMC) for 8 hours (Figure 1D). In contrast (and in accordance with its meiosis-specific function), *DMC1* expression is not induced after DNA damage treatment in WT plants. As expected, we observed strong induction of *DMC1* expression in all 3 independent *pRAD51::DMC1g* transgenic lines treated with 20 µM MMC for 8 hours (Figure 1D). We also tested expression of *HOP2*, *MND1* and *RAD54*, which are HR genes involved in the strand invasion process and known to be induced in somatic cells after treatment with DNA damaging agents [59–64]. Expression of all three genes was induced in both the transgenic plants and the wild-type controls (Figure 1D). Similar results were obtained when seedlings were treated with γ-rays (S2 Fig). Together, these data show that the *pRAD51::DMC1g* transgene allows high expression and induction of *DMC1* in both meiosis and mitosis and that this ectopic expression of *DMC1* does not appear to affect the expression of the other HR genes.

**Figure 1.**
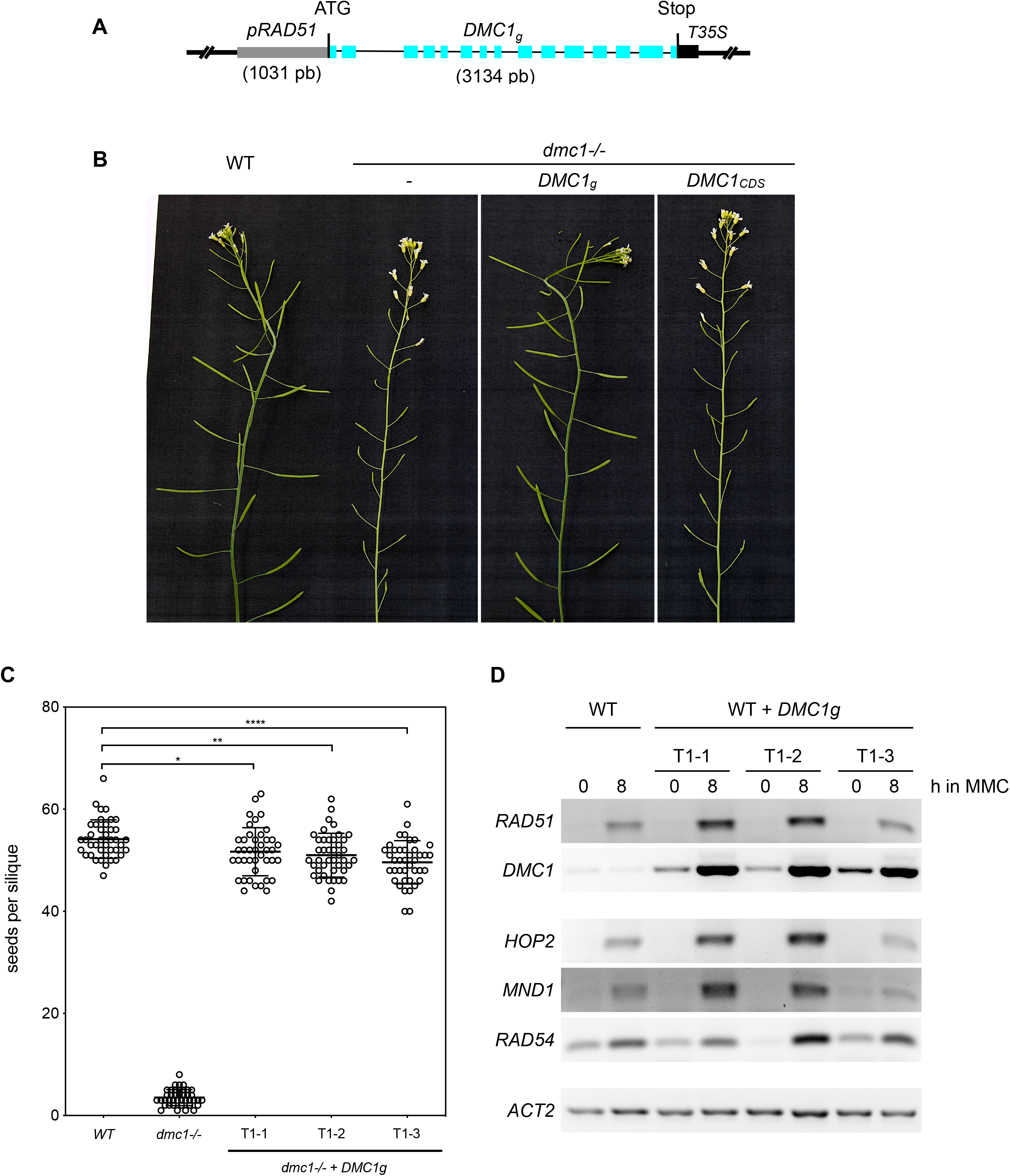
*pRAD51::DMC1g* restores fertility of the Arabidopsis *dmc1* mutant and induces expression of DMC1 in somatic cells. **(A)** Schematic representation of the *pRAD51::DMC1g* construct. *DMC1g* indicates *DMC1* genomic sequence. Exons are shown as blue rectangles. **(B)** Wild-type plants have long siliques full of seeds (mean ± S.D : 54.1 ± 0.6 seeds per silique), while *dmc1* mutants have very low fertility (mean ± S.D : 3.5 ± 0.25 seeds per silique). Expression of the *pRAD51::DMC1g* genomic sequence in *dmc1* mutants restores fertility. Mean ± S.D is shown; n = 4 plants, N = 10 siliques per plant. **(C)** number of seeds per silique in DMC1, *dmc1−/−,* and three *dmc1−/−* + *pRAD51::DMC1g* independent transformants (T1-1, T1-2, and T1-3) showing restored fertility. **(D)** RT-PCR expression analysis of *RAD51*, *DMC1*, *RAD54*, *HOP2* and *MND1* in 7-day-old, untreated and MMC-treated (8 h, 20 µM) seedlings expressing or not the *pRAD51::DMC1g* transgene. Expression was analyzed in wild-type plants and three independent transgenic lines (T1-1, T1-2, and T1-3). Actin is used as a loading control.

### Ectopic expression of DMC1 in somatic cells blocks RAD51-mediated DSB repair

We next analyzed whether RAD51-mediated DNA repair in somatic cells is affected by the presence of DMC1. We tested sensitivity of transgenic plants expressing *DMC1* in vegetative cells to the DNA damaging agent Mitomycin C (MMC). MMC forms DNA interstrand cross-link adducts, thereby producing DNA strand breaks that must be repaired by homologous recombination. In Arabidopsis, this is seen in the hypersensitivity of *rad51* mutants to MMC [39, 65]. Progeny of wild-type plants and *rad51* heterozygotes (*RAD51+/−*) carrying the *pRAD51::DMC1g* transgene were grown on solid media containing, or not, 20 µM Mitomycin C and growth was scored after 2 weeks (Figure 2 and S3 Fig). We tested the progeny of *RAD51+/−* heterozygous mutant plants as this permits analysis of the effect of vegetative expression of *DMC1* in both presence (*RAD51+/+* and *RAD51+/−*) or absence (*rad51−/−*) of RAD51. Plants grown under standard conditions did not exhibit any visible phenotypical differences indicating that ectopic expression of *DMC1* does not affect normal plant growth (Figure 2A). However, when grown in medium supplemented with MMC, transgenic plants showed strong hypersensitivity (Figure 2B and C), with 87% to 98% sensitive plants in the progeny of 3 independent *RAD51+/−* transformants (named T1-1, T1-2, and T1-3) carrying the *pRAD51::DMC1g* transgene (Figure 2B and C). This is comparable to the full sensitivity of *RAD51-GFP* plants (which behave as *rad51* mutants, [39, 66]) and in strong contrast to the 29% sensitive plants (corresponding to the segregating *rad51* homozygous mutants) observed for the progeny of *RAD51+/−* plants that do not carry the *pRAD51::DMC1g* transgene (Figure 2C). To confirm this, we selected *RAD51+/+* plants expressing the *pRAD51::DMC1g* transgene from the progeny of the three independent *RAD51+/−* transgenic lines tested above and examined their sensitivity to MMC. As expected, the three lines exhibited strong sensitivity (S3A-E Fig). Ectopic expression of *DMC1* in somatic cells thus has a dominant negative effect and we hypothesize that this occurs through precluding DSB repair mediated by RAD51.

**Figure 2.**
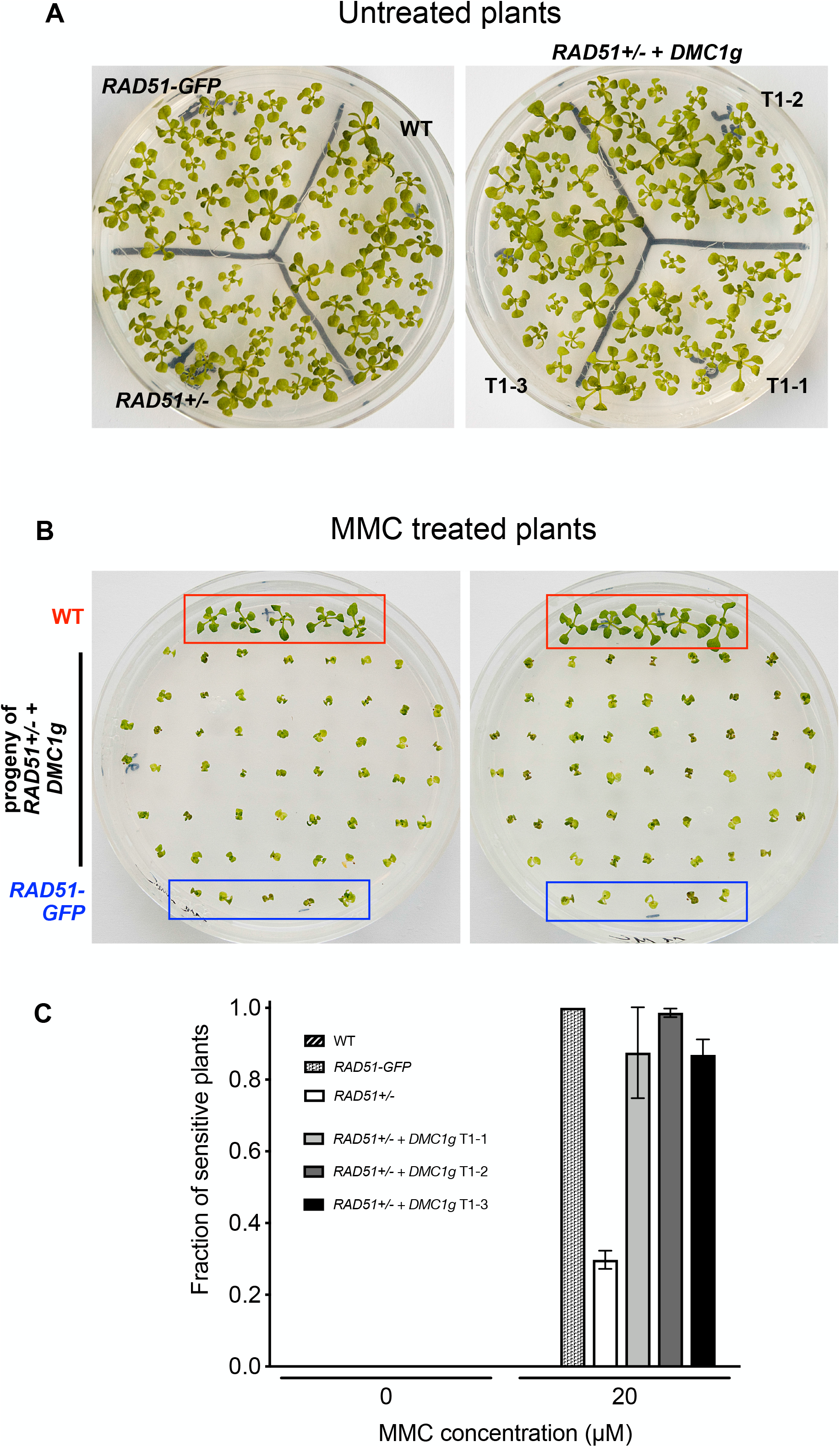
Mitomycin C hypersensitivity of transgenic seedlings expressing *pRAD51::DMC1g*. **(A)** No growth difference is observed between WT, *RAD51-GFP*, *RAD51+/−* heterozygous and *RAD51+/−* expressing *pRAD51::DMC1g* seedlings (two-week-old seedlings grown without MMC). **(B)** Pictures of two-week-old seedlings grown with 20 µM MMC. WT seedlings (depicted with red square) do not exhibit MMC sensitivity, in contrast to *RAD51-GFP* seedlings (blue square) and progeny of *RAD51+/−* transgenic lines expressing *pRAD51::DMC1g*. Pictures of progeny of two independent *RAD51+/−* transgenic lines expressing *pRAD51::DMC1g* are shown (T1-1 and T1-2). As control, WT (red square) and *RAD51-GFP* plants (blue square) were also grown on each plate. **(C)** Sensitivity of the seedlings was scored after 2 weeks and the fractions of sensitive plants (plants with less than 4 true leaves) are shown (3 biological repeats, each with N > 45 seedlings). Three independent *RAD51+/−* heterozygous lines expressing *pRAD51::DMC1g* (T1-1,T1-2 and T1- 3) were tested and all showed strong hypersensitivity to MMC.

### DMC1 does not substitute for RAD51 in DSB repair in somatic cells

An interesting observation from the above data is that DMC1 is not able to substitute for RAD51 in somatic cell DSB repair. Segregating *rad51* homozygotes are sensitive to MMC, even though they express *DMC1* (Figure 3B and C). This result might be surprising given the similar biochemical activities of the two proteins [16, 26, 32–34, 67]. Given that meiotic DMC1 activity requires the presence of a RAD51 nucleofilament [38, 39], a possible explanation for this is that if DMC1 also requires a RAD51 filament to be functional in somatic cells, then DMC1 will not repair DSB induced by MMC when expressed in the *rad51* mutant. To check this, we tested whether DMC1 would repair MMC-induced DSB when expressed in *RAD51-GFP* plants. The RAD51-GFP fusion protein forms a nucleofilament which supports the activity of DMC1 in meiosis but is not proficient for DSB repair in somatic cells [39]. All three *RAD51-GFP*_*pRAD51::DMC1g* independent lines tested were strongly sensitive to MMC (S3F-G Fig). Thus, in contrast to meiosis, the presence of the (inactive) RAD51-GFP filament is not sufficient to support DMC1 activity in somatic cells. This suggests that DMC1 activity requires the presence of meiosis-specific co-factors or, alternatively, that DMC1 activity is blocked in somatic cells by unknown factors.

**Figure 3.**
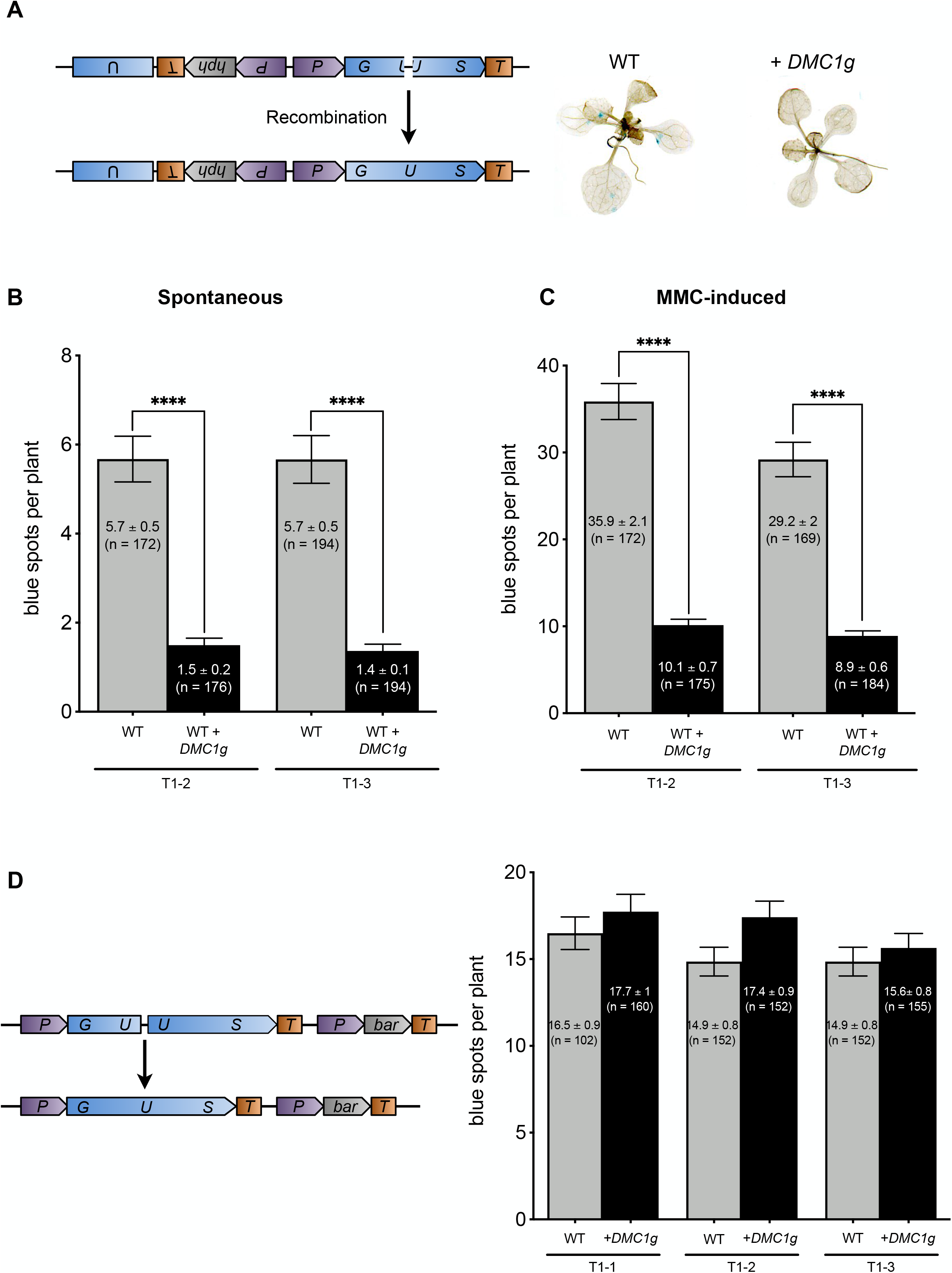
DMC1 inhibits RAD51-dependent somatic homologous recombination. **(A)** Schematic map of the *IU.GUS* chromosomal recombination reporter locus. Restoration of the functional *GUS* gene is visualized as blue spots in seedlings. Both spontaneous **(B)** and MMC induced **(C)** *IU.GUS* recombination rates are strongly reduced in wild-type seedlings expressing *pRAD51::DMC1g*. Mean numbers of GUS+ recombinant spots per plant ± standard errors of the means (SEM) are indicated for each genotype (n = 3, N > 45 seedlings). **** p < 0.0001 (Mann-Whitney test). The wild-type plants with and without *pRAD51::DMC1g* are sister plants from a transgenic line segregating for the *pRAD51::DMC1g* transgene. Analysis were performed using segregating plants from two independent transgenic lines (T1-2 and T1-3). **(D)** Schematic map of the *GU.US* chromosomal recombination assay and graph showing mean numbers of GUS+ recombinant spots per plant ± SEM. Three independent *GU-US* transformants expressing *pRAD51::DMC1g* (T1-1, T1-2, T1-3) and 2 WT (sister plants from a population segregating the *pRAD51::DMC1g* transgene) were analyzed (WT for T1-3 is the same as for T1-2). Three biological replicates, each containing at least 50 plants, were performed, except for the WT T1-1 for which only two replicates were performed. p > 0.05 (Mann-Whitney test).

### DMC1 specifically impairs RAD51-dependent recombination in somatic cells

Our results show that ectopic expression of *DMC1* impairs DSB repair in somatic cells and we hypothesize that this occurs through inhibition of RAD51 activity. However, another possible explanation for this hypersensitivity to DNA damage conferred by *DMC1* mitotic overexpression is that it might cause accumulation of nonproductive DMC1 nucleofilaments, which would block both RAD51-dependent and RAD51-independent recombinational repair, leading to genome instability and thus hypersensitivity to DNA damage. To check this, we tested the effect of ectopic expression of *DMC1* on somatic homologous recombination. If DNA damage hypersensitivity is caused by nonproductive DMC1 nucleofilaments, both RAD51-dependent and RAD51-independent recombination should be affected, while specifically blocking RAD51 should impact only RAD51-dependent recombination. We used the well-characterized *IU-GUS* recombination tester locus that specifically scores RAD51-dependent SDSA/GC recombination (Figure 3A) [68, 69]. This locus consists of a *β-glucuronidase* (*GUS*) gene with a 34-bp linker insertion, rendering it non-functional. Productive GUS recombination at the *IU.GUS* locus involves the use of an internal fragment of the *GUS* gene (5’ and 3’ deleted) placed in inverted orientation as donor sequence to restore the complete *GUS* gene (Figure 3A). The resulting functional *GUS* gene is scored histochemically as blue tissue sectors (Figure 3A). Spontaneous SDSA/GC recombination at *IU.GUS* was monitored in WT seedlings expressing or not *DMC1* (Figure 3B and S1 Data). We observed a dramatic reduction (4-fold) of SDSA/GC events in two independent transgenic lines (T1-1 and T1-2) compared to WT plants (Figure 3B and S1 Data). This reduction was confirmed after DNA damage induced by MMC treatment (Figure 3C and S1 Data). Seedlings were treated with 20 µM MMC and further grown for 3 days before monitoring recombination events. In wild-type, 30 to 35 SDSA/GC events/seedling were detected after treatment while seedlings ectopically expressing *DMC1* only showed 9-10 recombination events, corresponding to a 3.5-fold reduction (Figure 3C and S1 Data). Thus, ectopic expression of *DMC1* strongly impairs RAD51-dependent recombination.

To confirm that DMC1 specifically perturbs RAD51-dependent recombination, we also tested RAD51-independent recombination using the *DGU.US* recombination reporter locus (Figure 3D). This reporter consists of an I-SceI restriction site flanked by 3’ and 5’ truncated copies of the *GUS* gene in direct orientation and with a sequence homology overlap [68, 69]. Restoration of a functional *GUS* gene occurs through SSA recombination which is known to be RAD51-independent [66, 69]. As with the SDSA/GC assay, we measured SSA recombination efficiency in WT seedlings expressing or not *DMC1* (Figure 3D). Data from three independent lines (T1-1, T1-2, and T1-3) clearly showed that *DMC1* ectopic expression does not affect somatic *DGU.US* recombination (Figure 3D and S1 data). We concluded that somatic expression of DMC1 specifically impairs RAD51 activity, leading to inhibition of RAD51-mediated recombination and DSB repair in somatic cells.

### *DMC1* expression in somatic cells does not block RAD51 focus formation

We next tested whether expression of *DMC1* in somatic cells blocks RAD51 nucleofilament formation at DNA damage sites. The nucleofilament is the active molecular species that performs the homology search and strand exchange. Accordingly, defects in nucleofilament formation affect RAD51 activity. We analyzed RAD51 focus formation upon DNA damage in somatic cells as a proxy for RAD51 nucleofilament formation. We performed RAD51 immunofluorescence staining in root tip nuclei of 5-day-old seedlings treated or not with Mitomycin C (30 µM; Figure 4). As expected, no or very few foci were detected in root tip nuclei of non-treated seedlings (Figure 4A and D). In contrast, numerous RAD51 foci were detected in nuclei from root tips fixed 2 h or 8h after treatment with MMC (Figure 4B-D). In wild-type plants we observed an average of 2.3 foci (± 0.3; n=228) 2h after MMC treatment and this increases up to 15.8 foci (± 1.3; n=229) after 8h of MMC treatment (Figure 4 and S1 Data). Strikingly, similar distributions of RAD51 foci were observed in the root tips of two independent transgenic lines expressing *pRAD51::DMC1g* (T1-1 and T1-2; Figure 4B to D and S1 Data). Thus, *DMC1* expression in somatic cells does not affect RAD51 focus formation and thus probably RAD51 nucleofilament formation.

**Figure 4.**
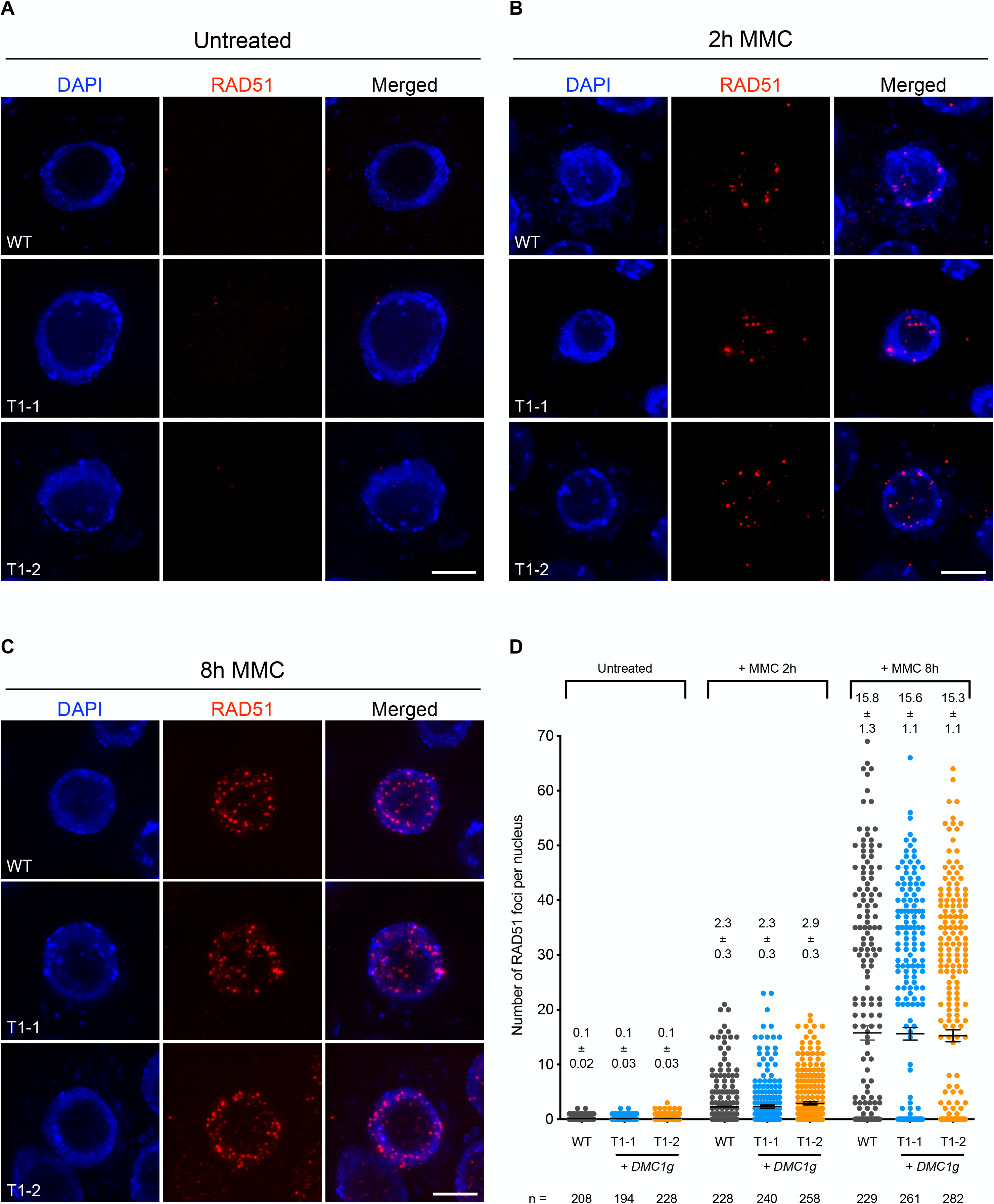
RAD51 foci in somatic cells of *DMC1* overexpressing seedlings. **(A-C)** Immunolocalization of RAD51 in root tip nuclei of wild-type plants expressing *pRAD51::DMC1g*, fixed just before **(A)**, and 2 **(B)** or 8 **(C)** hours after treatment with 30 µM MMC. T1-1 and T1-2 are two independent transgenic lines expressing *pRAD51::DMC1g*. MMC-induced RAD51 foci are clearly visible in nuclei of the treated plants. DNA is stained with DAPI (blue) and RAD51 foci (detected using an antibody against RAD51) are colored in red. Images are collapsed Z-stack projections of a deconvoluted 3D image stack. (Scale Bars : 5 µm). **(D)** Quantification of RAD51 foci in root tip nuclei of wild-type and two independent transgenic lines expressing *pRAD51::DMC1g* (T1-1 and T1-2) before and after MMC treatment. Means ± s.e.m are indicated for each genotype. More than 200 nuclei from at least 3 seedlings were analyzed per genotype.

### DMC1 focus formation in somatic cells upon DNA damage

Expression of *DMC1* in somatic cells impairs RAD51-mediated DSB repair and recombination, without blocking assembly of RAD51 filaments at DNA breaks. We thus hypothesized that DMC1 inhibits the strand invasion activity of the RAD51 nucleofilament in somatic cells. This conclusion prompted us to verify the ability of DMC1 to assemble at DNA breaks when expressed in somatic cells. We performed RAD51 and DMC1 immunofluorescence staining in root tip nuclei of 5-day-old seedlings treated with 30 µM Mitomycin C for 8 hours (Figure 5). As for RAD51, no DMC1 foci were detected without treatment, while numerous foci were observed after MMC treatment (Figure 5A and B). Interestingly, double immunolocalization of RAD51 and DMC1 shows that >90% of RAD51-positive cells were positive for DMC1 (n = 122 cells) and most, if not all, DMC1 positive cells were also positive for RAD51 (96%, n = 115 cells). We also observed strong overlap between RAD51 and DMC1 foci (Figure 5), with 65% of RAD51 foci overlapping with DMC1 foci and 82% of DMC1 foci overlapping with RAD51 foci (n = 221 foci analyzed). Scanning through overlapping regions in optical sections in several nuclei revealed that the signal intensities for RAD51 and DMC1 are very highly correlated (Fig 5C). This is confirmed by the Pearson correlation coefficient calculated from analysis of 100 randomly chosen foci (from 24 different cells), which ranges from 0.31 to 0.92 with a mean value of 0.71 (Figure 5D). RAD51 and DMC1 thus strongly co-localize, supporting the conclusion that they assemble together at DSB sites.

**Figure 5.**
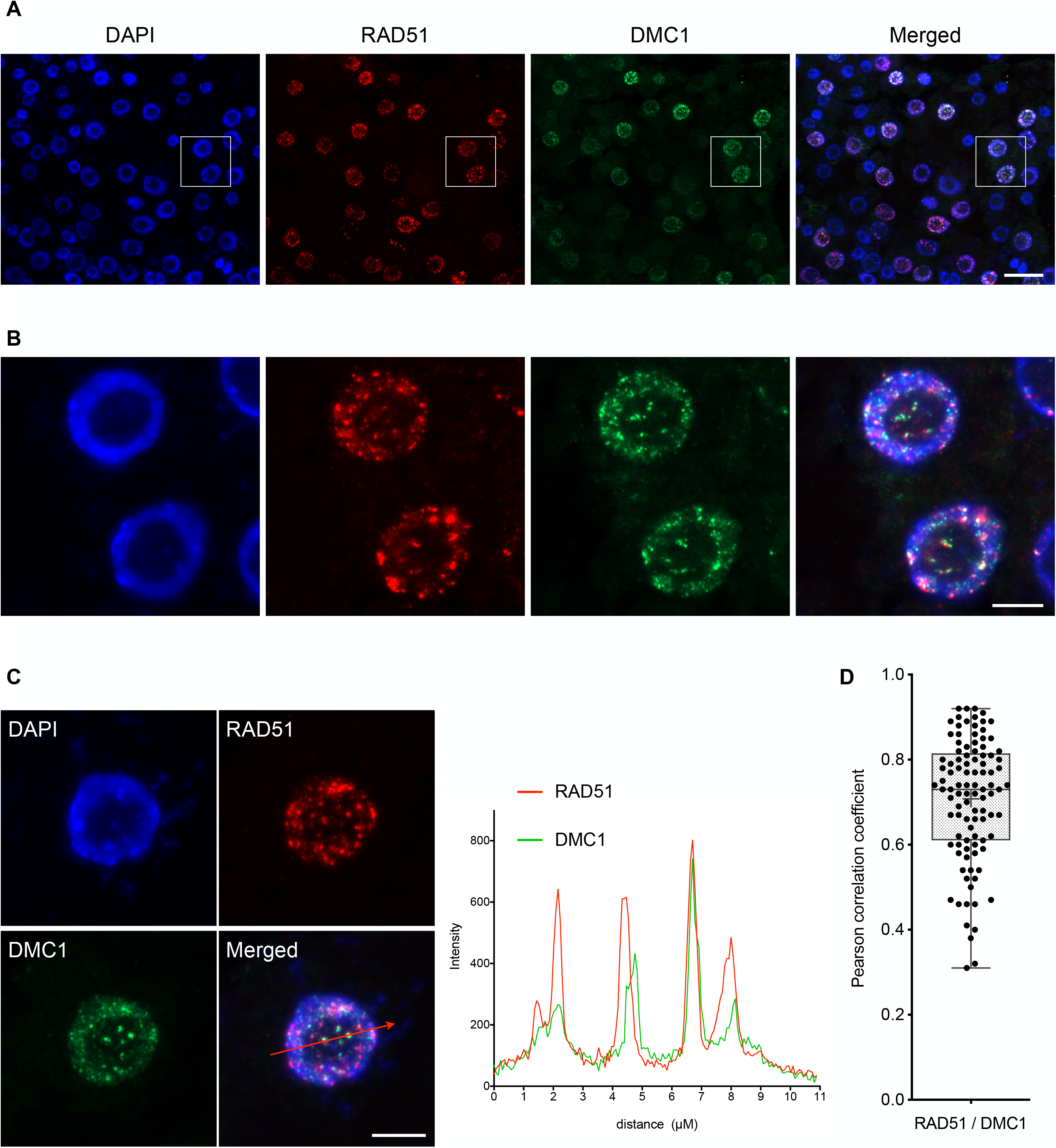
DMC1 foci are formed in somatic cells upon DNA damage. **(A-B)** Immunolocalization of RAD51 and DMC1 in root tip nuclei of wild-type plants expressing *pRAD51::DMC1g* fixed 8 hours after treatment with 30 µM Mitomycin C. **(B)** is a magnification of the area framed in **(A)**. DNA is stained with DAPI (blue), RAD51 foci are colored in red and DMC1 in green. Images are collapsed Z-stack projections of a deconvoluted 3D image stack. (Scale bars: 20 µm **(A)** and 5 µm **(B)**). **(C)** Example of RAD51 and DMC1 signal intensity distribution of the total amount of pixels through a section of the depicted stained nucleus (red line). RAD51 is shown in red and DMC1 in green. (Scale bar: 5 µm). **(D)** Range of Pearson correlation coefficients (PCCs) of RAD51/DMC1-positive foci formed after 8 h of MMC treatment. RAD51/DMC1 partially co-localized in foci with a mean value of PCCs = 0.63 (n = 40).

DMC1 can thus assemble at DNA breaks in somatic cells. The assembled DMC1 filament is non-productive but appears competent to block RAD51-dependent recombination.

### Downregulation of RAD51 recombinogenic activity requires DMC1 assembly at DSB sites

We next sought to test the capacity of non-productive DMC1 nucleofilaments to block RAD51-dependent recombination through expression of a mutant DMC1 protein, which forms nucleofilaments but is unable to catalyze DNA strand-transfer. We generated a *DMC1-II3A* variant with the rationale that mutating three conserved positively charged residues (R136, R308, and K318, S4 Fig) in the low-affinity DNA binding site (site II) of DMC1 will affect its recombinogenic activity without affecting ssDNA binding, as previously shown in yeast [38]. The DMC1 genomic coding sequence was synthesized with mutations to convert the three site II conserved residues (R136, R308, and K318, S4 Fig) to alanine. The Arabidopsis *DMC1-II3A* sequence was ten placed under the control of the *RAD51* promoter and the resulting construct was introduced in wild-type plants. Transgenic plants expressing *pRAD51::DMC1-II3A* were selected and presence of the protein examined (Figure 6A). Western blots confirmed that both the DMC1g and DMC1-II3A proteins are well translated in transgenic seedlings after Mitomycin C treatment (Figures 6A and S5). We then analyzed the presence of DMC1 foci in somatic cells through RAD51 and DMC1 immunofluorescence staining in root tip nuclei of 5-day-old seedlings treated with 30 µM Mitomycin C for 8 hours (Figure 6B). As seen above (Figure 5), numerous RAD51 and DMC1 foci are observed in transgenic seedlings expressing wild-type *DMC1g*. On the contrary, only RAD51 foci could be observed in transgenic plants expressing *pRAD51::DMC1-II3A* (Figure 6B). The inability of our DMC1-II3A to form foci was then confirmed in meiotic cells. The DMC1-II3A sequence was placed under the control of the *DMC1* promoter and the construct was introduced into *DMC1/dmc1* heterozygous plants. Homozygous *dmc1/dmc1* transformants expressing the *DMC1-II3A* transgene were selected. As anticipated, all 13 *dmc1/dmc1* mutant plants expressing the *DMC1-II3A* that we obtained were almost sterile. Counting the number of seeds per silique in two independent transformants revealed that these plants exhibit a *dmc1-like* phenotype (not the expected *rad51*-like fragmentation phenotype), with 2-3 seeds per silique (S6 Fig). Interestingly, we also observed a strong dominant negative phenotype in WT plants showing strongly reduced fertility (S6 Fig). Immunolocalization confirmed the absence of DMC1 foci in *dmc1* mutants expressing *DMC1-II3A* (S7 Fig). Accordingly, immunolocalization of the synaptonemal complex transverse filament protein ZYP1 revealed defective synapsis (*dmc1* phenotype) in these plants (S7 Fig). Thus, in contrast to the analogous protein in budding yeast [38], the Arabidopsis DMC1-II3A does not appear to stably associate and/or polymerize on ssDNA and thus does not support DMC1 presynaptic nucleofilament formation.

**Figure 6.**
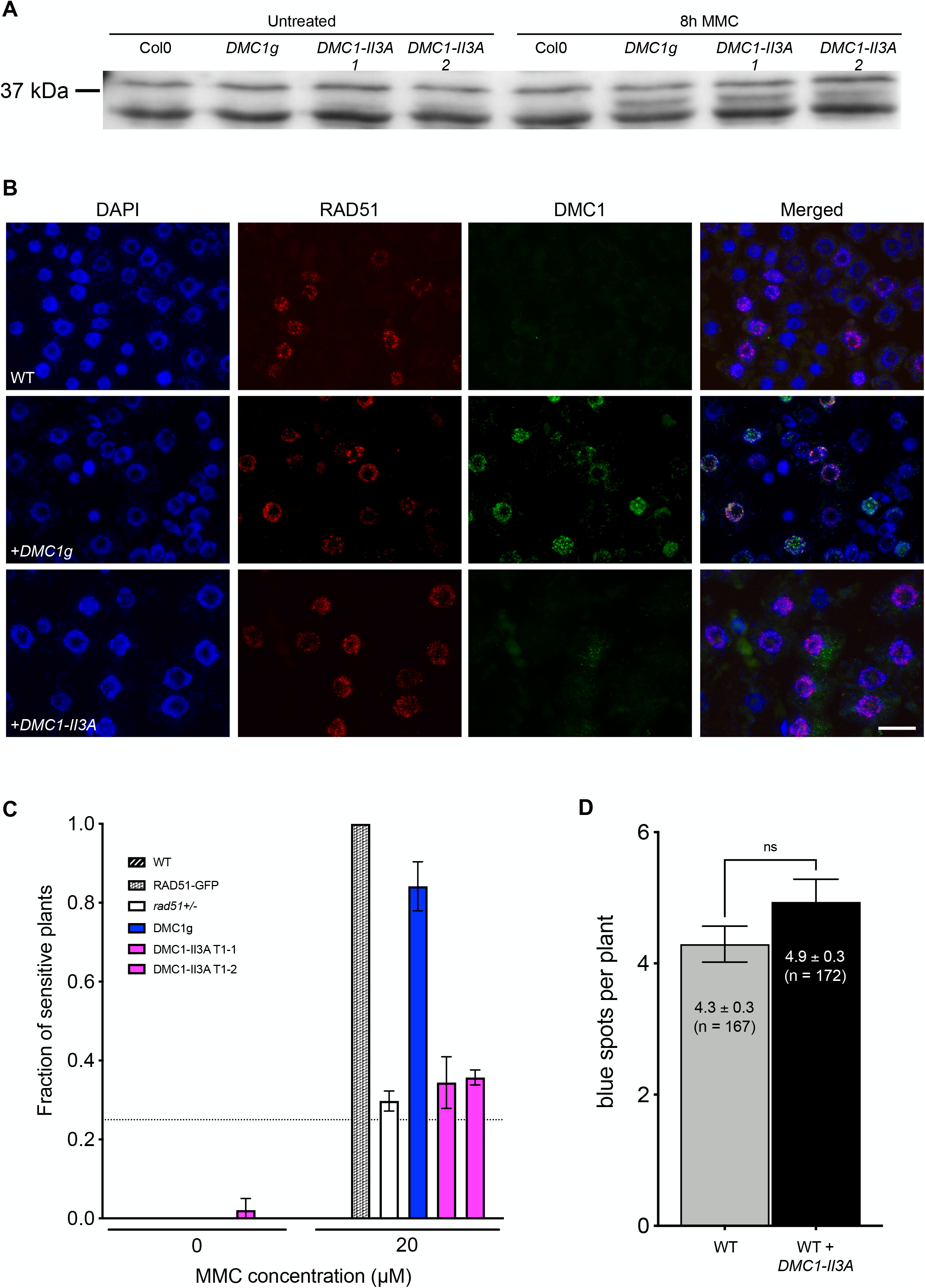
DMC1-II3A variant does not form focus nor prevent RAD51-mediated recombination in somatic cells. **(A)** DMC1g and DMC1-II3A protein are induced by MMC treatment. Total proteins were extracted from 1-week-old seedlings treated or not with 30 µM MMC for 8 hours and DMC1 abundance measured. No DMC1-specific band (37 kDa) is observed in untreated plants in both wild-type and transgenic plants. In contrast, while DMC1 is still absent in wild-type plants after DNA damage treatment, it becomes clearly visible in transgenic plants expressing either *DMC1g* or *DMC1-II3A*. **(B)** Immunolocalization of RAD51 and DMC1 in root tip nuclei of wild-type plants, transgenic plants expressing *pRAD51::DMC1g* or *pRAD51::DMC1-II3A* fixed 8 hours after treatment with 30 µM Mitomycin C. DNA is counterstained with DAPI (blue). Images are collapsed Z-stack projections of a deconvoluted 3D image stack. (Scale bar : 20 µm). **(C)** Expression of DMC1-II3A does not render plant hypersensitive to MMC nor **(D)** prevent RAD51-dependent somatic homologous recombination. **(C)** Sensitivity of seedlings was scored after 2 weeks of growth with 20 µM MMC and the fractions of sensitive plants (plants with less than 4 true leaves) are shown (3 biological repeats, each with N > 45 seedlings). Two independent transgenic lines expressing *pRAD51::DMC1-II3A* (T1-1 and T1-2) were tested and none exhibit MMC hypersensitivity. **(D)** Mean numbers of GUS+ recombinant spots per plant ± standard errors of the means (SEM) are indicated for each genotype. (p < 0.05; Mann-Whitney test). The wild-type plants with and without *pRAD51::DMC1-II3A* are sister plants from a transgenic line segregating for the *pRAD51::DMC1-II3A* transgene.

We tested impacts of DMC1-II3A on MMC hypersensitivity and RAD51-mediated recombination using the *IU-GUS* recombination tester locus described above. As expected by the absence of DMC1 foci, transgenic seedlings expressing the *DMC1-II3A* transgene exhibit neither MMC hypersensitivity (Figure 6C) nor impairment of RAD51-dependent recombination (Figure 6D and S1 Data). The absence of mitotic phenotype of this non-nucleofilament forming mutant DMC1-II3A protein concords with DMC1-mediated downregulation of RAD51 recombinogenic activity through DMC1 assembly at DSB sites.

### Expression of dominant negative DMC1-GFP blocks RAD51 activity in meiosis

Expression of a nucleofilament-forming DMC1 in mitotic cells blocks RAD51 activity, in accord with the model of inhibition of RAD51 recombinogenic activity by the DMC1 nucleofilament. In meiotic cells, blocking RAD51-mediated recombination in the absence of DMC1-mediated recombination should lead to meiotic chromosomal fragmentation, in contrast to the intact univalent chromosomes observed in the *dmc1* null mutant. Unfortunately, the DMC1-II3A protein cannot be used to test the model as it appears not to form nucleofilaments in Arabidopsis. We thus generated a *DMC1-GFP* translational fusion (that we should be able to visualize) with the rationale that the DMC1-GFP protein will be inactive for DNA strand-transfer, while retaining the capacity to form nucleofilaments (analogously to RAD51-GFP in Arabidopsis, mammals and yeast [39, 41, 70–72] and DMC1-GFP in fission yeast [73]).

We amplified the DMC1 genomic coding sequence (including introns but without the stop codon) and 2.7 kb of upstream sequence from DNA of wild-type Arabidopsis and fused the *GFP* coding sequence to the 3’ end of the DMC1 open reading frame. The fusion construct was introduced into *DMC1/dmc1* heterozygous plants and transformants expressing the *DMC1-GFP* translational fusion protein were selected. As anticipated, all 8 *dmc1/dmc1* plants expressing the *DMC1-GFP* that we obtained were sterile, confirming that the fusion protein is not able to complement meiosis of the Arabidopsis d*mc1* mutant and is thus not functional (S8 Fig). Analogously to the RAD51-GFP in Arabidopsis, DMC1-GFP acts as a dominant negative, with more than 80% of wild-type plants (WT) expressing DMC1-GFP (59 of 71) being sterile or showed strongly reduced fertility (S8 Fig).

To directly verify whether the DMC1-GFP fusion protein interferes with RAD51 activity we next investigated male meiosis by staining chromosomes with 4’,6-diamidino-2-phenylindole (DAPI) (Figure 7). Presence of the *DMC1-GFP* leads to clear chromosome fragmentation (*rad51*-like phenotype) in *DMC1-GFP dmc1−/− RAD51+/−* (compare with *dmc1−/− RAD51+/−* alone in which no fragmentation is observed; Figure 7B). Fragmentation was also observed in *DMC1-GFP DMC1+/+ RAD51+/−* plants (Figure 7A and B). The presence of the DMC1-GFP protein thus blocks meiotic RAD51 activity. We note that the penetrance of this phenotype is incomplete and appears to depend upon the presence of wild-type DMC1 protein as well as being impacted by the dosage of RAD51. In contrast to *DMC1-GFP dmc1−/− RAD51+/−, DMC1-GFP dmc1−/− RAD51+/+* mutants show a *dmc1-like* phenotype with impaired pairing and synapsis, intact Metaphase I univalents and random chromosome segregation during Anaphase I (Figure 7A and B). Likewise, *DMC1-GFP DMC1+/+ RAD51+/+* plants also have a *dmc1-like* phenotype (Figure 7 and S8 Fig).

**Figure 7.**
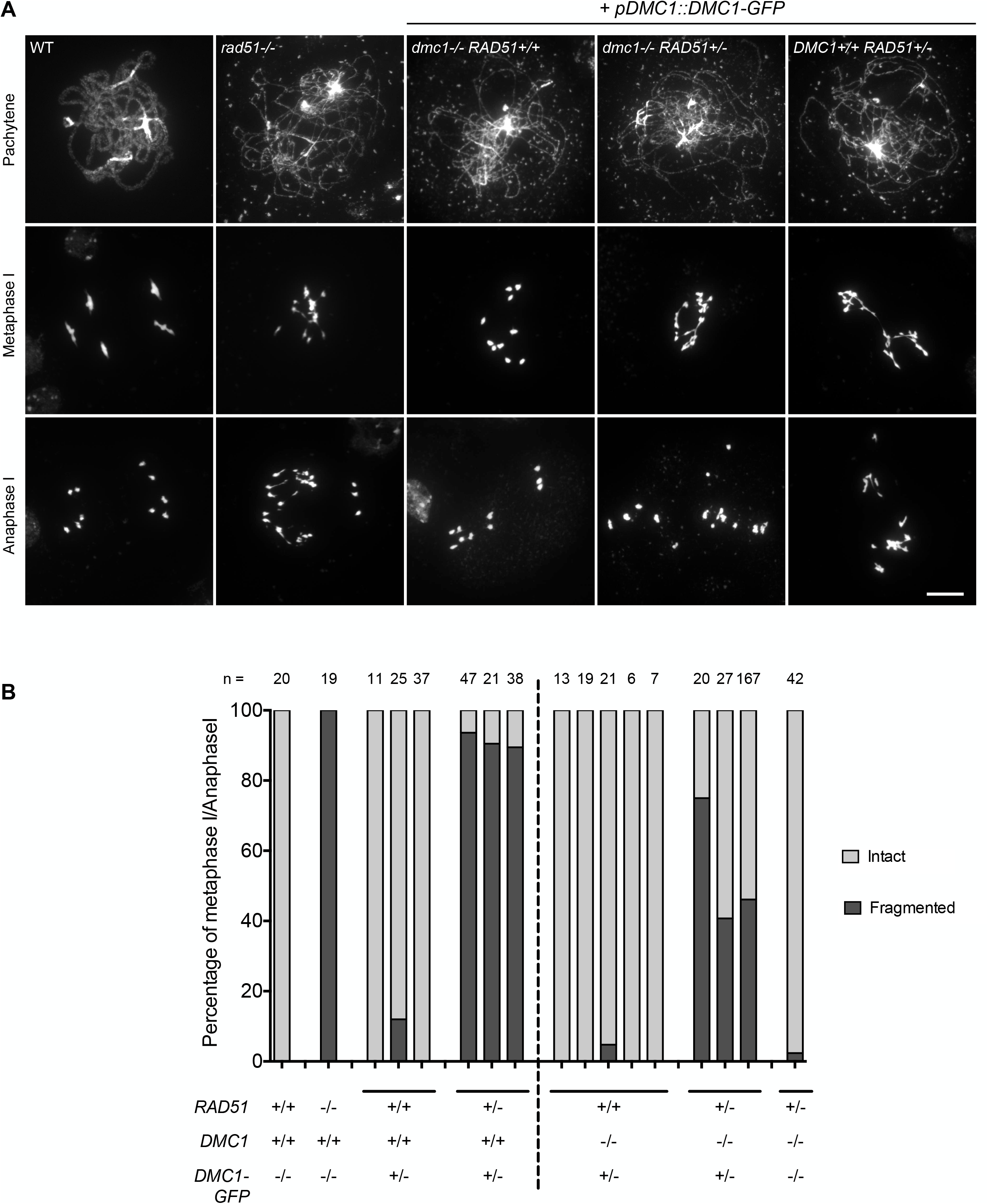
Expression of DMC1-GFP fusion protein inhibits RAD51-mediated repair of meiotic DBS. **(A)** DAPI staining of chromosomes during meiosis. Wild-type cells show pairing and synapsis of homologous chromosomes in pachytene, five bivalents at metaphase I and two groups of five chromosomes at anaphase I. *rad51* mutants exhibit defective synapsis and extensive chromosome fragmentation. Meiotic spreads in *dmc1−/− RAD51+/+* plants expressing *DMC1-GFP* revealed a *dmc1-like* phenotype with defective synapsis but ten intact univalents at metaphase I, which randomly segregate at anaphase I. In contrast, extensive fragmentation is observed in both *dmc1−/− RAD51+/−* and *DMC1+/+ RAD51+/−* plants expressing *DMC1- GFP*. (Scale Bar: 10 µm). **(B)** Quantification of chromosome fragmentation. The genotype of the plants is indicated below each bar and number of cells analyzed is listed above.

These analyses were extended through immunolocalization of DMC1 to examine the presence of DMC1-GFP foci in meiotic cells (S9 Fig). Using a DMC1-specific antibody, numerous DMC1 foci were observed in wild-type plants expressing DMC1-GFP while none were detected in *dmc1−/−* expressing *DMC1-GFP* (S9 Fig). Yet, the phenotype observed in the DMC1-GFP transgenic plants indicates that the fusion protein is well-expressed and this is supported by observation of GFP fluorescence in nucleus of live meiocytes of wild-type and *dmc1* plants expressing *DMC1-GFP* (S9 Fig). We thus conclude that DMC1-GFP is well-expressed and its nucleofilament-forming ability depends upon the presence of WT DMC1 protein, or alternatively that it impedes DMC1 nucleofilament formation. Thus, in DMC1 + DMC1-GFP meiosis, the DMC1-GFP-containing nucleofilaments block RAD51 activity (*rad51*-like phenotype), while no DMC1-GFP nucleofilaments form in the absence of DMC1 and this results in a *dmc1-like* phenotype. These effects on the penetrance of the DMC1-GFP phenotype point both to a particularity of the DMC1-GFP protein and to the importance of a tightly controlled stoichiometry of both recombinases in meiosis, but do not affect the conclusion that DMC1-GFP impedes DMC1-mediated recombination and triggers downregulation of meiotic RAD51-mediated recombination.

## Discussion

### DMC1 potentially prevents RAD51-mediated recombination

The data presented here support the argument that DMC1 plays a key role in preventing the activity of RAD51 in driving meiotic recombination in Arabidopsis. We present our conclusions in the context of a model on the formation and activity of RAD51 and DMC1 nucleofilaments in Figure 8. In most studied eukaryotes, efficient meiotic recombination requires the presence of both RAD51 and DMC1 [16] (Figure 8A). In our model, RAD51 and DMC1 loading is represented according to the symmetrical loading model with RAD51 and DMC1 homotypic filaments on both DSB ends of a break. This is the most recently supported model and has been shown to better explain regulation of RAD51 and DMC1 activity [16, 17, 21–24, 26, 27]. We note however that asymmetric loading has also been previously suggested (in particular in Arabidopsis) and is thus also possible [19]. The nucleofilament is then the active molecular species and data from budding yeast, mouse and Arabidopsis have demonstrated that DMC1 is the active recombinase during meiosis with RAD51 being an essential co-factor for DMC1, but its strand exchange activity being inhibited [21, 38–40, 46, 47, 57, 74] (Figure 8A). Thus, absence of RAD51 leads to strong defects in DMC1 loading and therefore defects in meiotic DSB repair [19, 38, 58, 75, 76] (Figure 8B). In budding yeast, down-regulation of Rad51 is achieved by at least two mechanisms involving the meiosis-specific kinase Mek1 : (i) Mek1 phosphorylates and stabilizes the meiosis-specific Hed1 which competes with Rad54 for binding to Rad51 [45, 49, 51, 52], and (ii) Mek1 phosphorylates Rad54, reducing its affinity for Rad51 and thereby its activity [50]. However, homologs of Mek1 and Hed1 have not been identified, nor has the mechanism by which RAD51 is downregulated been described in organisms other than budding yeast. We recently confirmed that Arabidopsis RAD54 is essential for RAD51-mediated recombination and that this includes meiotic DSB repair in *dmc1* mutant plants [74] (Figure 8C and E). This study thus further emphasized that the RAD51-RAD54 pathway is down-regulated or plays a minor role in wild-type Arabidopsis meiosis.

**Figure 8.**
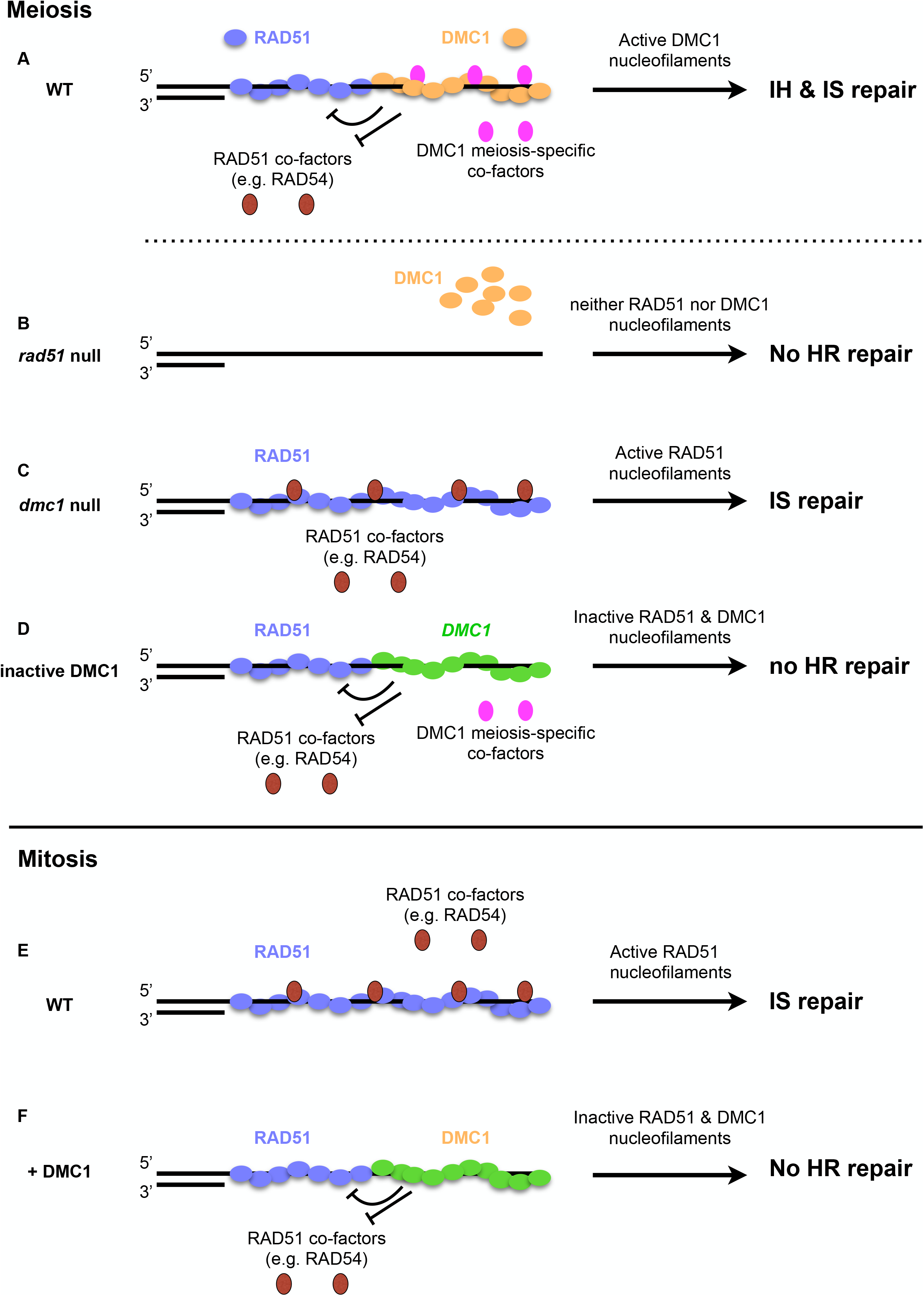
Hypothetical model for the activity of RAD51 and DMC1 nucleofilaments. **(A)** During meiotic recombination, RAD51 and DMC1 polymerize on ssDNA to form nucleofilaments. DMC1 is the active nucleofilament but requires the presence of RAD51 nucleofilaments and additional co-factors. Presence of the DMC1 nucleofilament inactivates the RAD51 nucleofilament either through an unknown signaling pathway, through inhibiting interaction with co-factors or through nucleofilament remodeling. **(B)** In absence of RAD51, DMC1 does not form nucleofilaments and DSB repair is defective. **(C)** Absence of DMC1 allows normal polymerization of RAD51 and interaction with co-factors. RAD51 nucleofilaments are thus proficient for DSB repair using sister chromatid templates. This is reminiscent of the situation in mitosis **(E)** where RAD51 is the sole recombinase. Its polymerization and interaction with co-factors such as RAD54 lead to an active nucleofilament proficient for DSB repair**. (D and F)** Addition of non-functional DMC1 (e.g. *DMC1-GFP*) in **(D)** meiosis or **(F)** mitosis, leads to RAD51 down-regulation. In mitosis, DMC1 may be kept inactive presumably through absence of meiosis-specific co-factors or chromosomal features. We note that the spatial organization of recombinases and accessory factors depicted in the model is hypothetical and thus other organizations are possible (reviewer in [16, 17, 25]). In particular, although the symmetrical loading of RAD51 and DMC1 is currently favored, it has been formally demonstrated in yeast only and it is not yet clear whether it applies for other organisms. This is particularly true for Arabidopsis where asymmetric loading has previously been suggested [19].

This work in Arabidopsis suggests that meiotic RAD51-downregulation is mediated by DMC1 (Figure 8D and F). In Arabidopsis, previous reports have already suggested that DMC1 may inhibit RAD51, based on observations that deletion of DMC1 in some mutant backgrounds leads to activation of RAD51 [56–58, 74] (Figure 8C). In Arabidopsis *hop2* or *mnd1* mutants, extensive DNA fragmentation occurs in meiosis as a result of defective recombination and absence of DMC1 in these mutants alleviates this DNA fragmentation, presumably through reactivation of RAD51 [57, 58]. That DMC1 is able to inhibit RAD51 recombinogenic activity has also been suggested in budding yeast [18, 46]. Dmc1 is known to favor inter-homolog recombination while Rad51 favors sister chromatid recombination. Accordingly, *hed1* mutants exhibit defects in homolog bias as a result of Rad51 activation, and this effect is stronger in the *dmc1 hed1* double mutant than in the *hed1* single mutant [18, 46] suggesting that Rad51 is even more active in absence of Dmc1. Similar conclusions have been drawn from comparisons of *mnd1 hed1* and *dmc1 hed1* mutants [46]. In the *mnd1* mutant, both DMC1 and RAD51 are present but strand-exchange is inhibited, resulting in meiotic arrest, unrepaired DSB and the virtual absence of crossovers. These defects are alleviated by mutating HED1, but to a lesser extent than in *dmc1 hed1* double, and *mnd1 dmc1 hed1* triple mutants [46]. As in Arabidopsis, this is however indirect evidence and whether this DMC1 inhibitory effect occurs in a wild-type context or only in certain mutant backgrounds is not clear.

We have expressed a non-functional *DMC1-GFP* allele in meiosis and DMC1 in somatic cells and show that they block RAD51 activity, strongly supporting the hypothesis that DMC1 has the potential to prevent RAD51-mediated recombination (Figure 8D and F). We note however that whether this DMC1-dependent down-regulation of RAD51 also holds true in WT meiotic cells is indirectly inferred here through our data in somatic cells (Figure 8F), the meiotic phenotypes of several mutants described above, and the meiotic phenotype of the DMC1-GFP (Figure 8D). The dominant negative nature of the DMC1-GFP on DMC1 does not allow us to unequivocally conclude whether the meiotic RAD51 inhibition is the result of an intrinsic activity of DMC1 itself, or whether it comes from the presence of the GFP domain. We sought to directly test this though expression of a non-recombinogenic *DMC1-II3A* allele in meiosis but unfortunately failed to do so due to the inability of Arabidopsis DMC1-II3A to form DMC1 foci. We do note however, that this inability of the Arabidopsis protein to form foci is in striking contrast with budding yeast Dmc1-II3A and underlines the existence of differences in behavior and/or regulation of DMC1 between these species.

### Mechanistic insights into DMC1-dependent down-regulation of RAD51 recombinogenic activity

How DMC1 could prevent RAD51 activity is intriguing since RAD51 is an essential co-factor for DMC1 and DMC1 must thus couple RAD51 inhibition with the supporting role of RAD51 on its own activity. Previous work has shown that the meiotic nucleofilament-forming ability, not the strand-exchange activity of RAD51 is needed for supporting DMC1 activity in meiosis and that meiotic RAD51 nucleofilaments form normally in the absence of DMC1 [19, 38–40, 58, 74, 76, 77].

RAD51 recombination can be downregulated on several levels by affecting various RAD51 activities. This includes RAD51 loading on RPA-coated ssDNA, stabilization of the nascent nucleoprotein filament by controlling its assembly/disassembly, RAD51 ATPase activity, reduction of RAD51’s capacity to bind dsDNA or even remodeling of the RAD51 nucleoprotein filament into a conformation that is optimally permissive for DNA strand exchange. The work presented here confirms that somatic RAD51 focus formation is not impaired by ectopic expression of *DMC1*, even though the presence of DMC1 blocks their activity in recombination. Thus, the prevention of RAD51-dependent recombination by DMC1 is mediated through down-regulation of activities of RAD51 nucleofilaments, not of their formation. This interpretation is supported by (i) the presence of inactive DNA damage-induced DMC1 foci in somatic cells, (ii) the down-regulation of RAD51 DSB repair in meiosis by a non-functional dominant negative *DMC1-GFP* allele and (iii) it concords with the characterization of several Arabidopsis null mutants (e.g. *hop2* and *mnd1*), in which meiotic DSB repair defects lead to chromosome fragmentation even though both RAD51 and DMC1 form foci at wild-type levels [56-58, 78, 79]. The Arabidopsis *sds* mutant (a meiosis-specific cyclin-like protein) has a *dmc1-like* meiotic phenotype with an absence of DMC1 focus formation. Meiotic DSB are however fully repaired by RAD51, most probably using sister chromatid templates [56, 79–82]. Interestingly, deletion of the AAA-ATPase FIDGETIN-LIKE-1 (FIGL1) or its partner FLIP partially restores DMC1 focus formation in *sds* mutants but this is associated with important chromosome fragmentation. That this is linked to the presence of DMC1 foci is confirmed by the presence of intact chromosomes in a triple *sds figl1 dmc1* mutant [56, 79].

Remarkably, RAD51 is not the only protein inhibited by DMC1. In budding yeast, DMC1 is able to specifically inhibit the ATPase activity (hence the motor activity) of the antirecombinase Srs2, even though Srs2 retains its ability to bind DMC1-ssDNA filaments [83]. The RAD51-GFP protein that is proficient for supporting DMC1 in meiotic recombination exhibits normal ATP binding and hydrolysis *in vitro* arguing against regulation of RAD51 ATPase activity [41].

Several other non-exclusive scenarios can be envisioned. For instance, DMC1 may inhibit RAD51 through effects on RAD51’s interaction with essential co-factors. It is tempting to speculate that one such co-factor could be RAD54, which could be functionally analogous to the RAD51-RAD54 regulation of budding yeast (Figure 10). Arabidopsis RAD54 is required for RAD51 function in somatic cells [63, 84–86] and we have recently shown that it is also essential for RAD51 recombinogenic activity in meiotic cells in absence of DMC1 [74].

Another possibility is that DMC1 affects the properties of the RAD51 nucleofilament. The nucleofilament is the active RAD51 species and filaments with distinct properties have been described. Super-resolution microscopy, single molecule imaging and chromatin immunoprecipitation approaches have revealed that RAD51 and DMC1 form separate homotypic filaments on both ends of a DSB [21–24]. Furthermore, meiotic DMC1 function depends upon the presence of RAD51 nucleofilaments (not their strand-invasion activity) and RAD51 foci form normally in *dmc1* mutants of Arabidopsis, yeast and mice, while numbers of DMC1 foci are significantly reduced in *rad51* mutants [19, 38, 58, 76, 77]. Thus, DMC1 may cap RAD51 filaments and block their activity. In agreement with this, our results show strong overlap of RAD51 and DMC1 foci in somatic cells.

Additionally, DMC1 may change the conformation of the RAD51 filaments. RAD51 nucleofilaments with different conformational states have been described by crystallography : one with an “extended” conformation and another with a “condensed” conformation [87]. The “extended” conformation is presumed to be the open, active state of the filament and the “condensed” conformation to represent a closed state of the filament. Dynamic arrangement of the nucleofilament is central to its function and it is thus possible that binding of DMC1 blocks RAD51 in a closed state.

An interesting additional conclusion from our data is that DMC1-dependent down-regulation of RAD51 requires a controlled stoichiometry of the two recombinases. Expression of the dominant negative *DMC1-GFP* allele in meiosis inhibits RAD51 repair only in a *RAD51+/−* (not *RAD51+/+)* context. This suggests that RAD51 down-regulation requires sufficient amounts of DMC1 and presumably DMC1-nucleofilaments. In accordance we found that inhibition of RAD51 activity was stronger in presence of both DMC1-GFP and endogenous DMC1, than by DMC1-GFP alone (compare fragmentation *in DMC1-GFP DMC1+/+ RAD51+/−* vs. *DMC1-GFP dmc1−/− RAD51+/−* plants in Figure 6). We speculate that DMC1-GFP alone does not form sufficient stable and/or abundant filaments while mixed DMC1 endogenous/DMC1-GFP filaments are more stable and/or abundant and thus more prone to inhibit RAD51. In support of this, we observed DMC1 foci in *DMC1-GFP DMC1+/+* but not in *DMC1-GFP dmc1−/−* plants, underlining the importance of tight regulation in meiotic cells allowing sufficient and timely expression of *DMC1* versus *RAD51*.

### Replacement of RAD51 in somatic cells with DMC1 : mitotic HR versus meiotic HR

To investigate whether DMC1 inhibits RAD51, we took the approach of ectopically expressing *DMC1* in somatic cells in which normally only RAD51 is present. RAD51 and DMC1 share similar biochemical properties and catalyze essentially the same DNA strand exchange reaction [16, 26, 33, 34, 67], although DMC1 fibres are able to accommodate unpaired base mismatches [88–92] and results from fission yeast indicate opposite polarities of nucleofilament polymerization of the two proteins [16, 93]. It is not yet clear whether functional differences between RAD51 and DMC1 result from distinct biochemical properties or influence of different co-factors. Our data may provide new insights into this important question. First, immunolocalization studies showed that DMC1 can form DNA damage-induced foci in somatic nuclei, suggesting that DMC1 assembly on ssDNA does not require meiosis-specific co-factors. Although recruitment of DMC1 to sites of DNA breaks requires meiosis-specific co-factors in budding yeast, such as Mei5-Sae3 complex, no such factors have been identified in Arabidopsis and our data suggest that DMC1 may not require any such factors for loading at DNA breaks. We further show that expression of *DMC1* rescues neither somatic HR, nor DNA damage hypersensitivity of *rad51* mutants. Thus, notwithstanding its ability to form nucleofilaments, DMC1 is not able to replace RAD51 for repair in somatic cells. This indicates either that DMC1 strand-exchange activity does require meiosis-specific co-factors and/or chromosomal features or alternatively that DMC1 strand-exchange is inhibited in somatic cells by some yet unknown factors. One evident candidate meiosis-specific co-factor would be the HOP2/MND1 complex, an essential co-factor for DMC1 in meiosis [57, 58, 94–97]. This would fit with an earlier study on the different meiotic phenotypes of the null and hypomorphic Arabidopsis *hop2* mutants, which proposed a model in which transient inhibition of RAD51 by DMC1 is alleviated by interaction between DMC1 and the HOP2/MND1 complex. However, a role for HOP2/MND1 in mitotic RAD51-mediated recombination has been described in mammals and Arabidopsis [58, 62, 98, 99]. HOP2/MND1 expression is strongly responsive to DNA damage in Arabidopsis somatic cells and Arabidopsis *mnd1* mutants have also been shown to be sensitive to DNA damage [59–64]. HOP2/MND1 is thus present and functional in somatic cells and is unlikely to be the “missing” meiotic DMC1 co-factor. In this context, our system also offers a unique opportunity to investigate RAD51-mediated HR vs. DMC1-mediated HR. It would be interesting to see how adding known meiotic specific factors (e.g. REC8,…) to the DMC1 reaction affects somatic recombination. This would provide important information concerning the evolution of the meiotic program and the specificities provided by DMC1 [100].

## Materials and Methods

### Plant Material and Growth Conditions

The lines used in this study were *rad51-1* [54], *dmc1-1* [53] and *RAD51-GFP* [39]. Seeds were stratified in water at 4°C for 2 days and grown on soil or *in vitro* on 0.8% agar plates with 1% sucrose and half-strength Murashige and Skoog salts medium (M0255; Duchefa Biochemie). Plants were then cultivated in a growth chamber with a 16/8-hour light/dark cycle, temperature 23°C and 60% relative humidity.

### Cloning of DMC1 and plant transformation

For expression of *DMC1* in vegetative cells (here called *DMC1g*), the complete genomic region from the ATG to the stop codon (2793 bp) of *DMC1* was amplified (forward primer ATGATGGCTTCTCTTAAGTAAGT and reverse primer CTAATCCTTCGCGTCAGC), inserted into pDONR-Zeo and verified by sequencing. The fragment was then cloned into a GATEWAY destination vector pMDC32 [101] in which the *35S* promoter was replaced by the Arabidopsis *RAD51* promoter as described in [39].

For *DMC1-II3A*, the complete genomic region from the ATG to the stop codon of *DMC1* (as above) was synthesized with mutation to convert R136, R308 and K318 into Alanine (S4 Fig). For expression in meiosis, the synthesized *DMC1-II3A* was cloned into the GATEWAY destination vector pMDC32 in which the 35S promoter was replaced by the DMC1 promoter (2.7kb 5’ upstream sequence of *DMC1*) using an AscI/SbfI digest. The DMC1 promoter was amplified from *Landsberg erecta* (*Ler-0*) DNA using forward primer GCGTGCCTGCAGGCACACCTAATCGGTGATTGCC and reverse primer GCTTGGCGCGCCtttctcgctctaagagtctctaa).

For translational GFP fusions, the DMC1 genomic region without stop codon and a 2.7kb 5’ upstream sequence of *DMC1* was amplified from Landsberg DNA (forward primer CCACACCTAATCGGTGATTG and reverse primer ATCCTTCGCGTCAGCAATG) and cloned into a GATEWAY destination vector using the strategy described in [39].

Plasmids were then inserted in an *Agrobacterium tumefaciens* C58C1 strain, which was subsequently used to transform Arabidopsis plants by the floral dip method [102]. Seeds from the Agrobacterium-treated plants were sown on soil and transformants were selected for Hygromycin or Basta resistance.

### Semi-quantitative RT-PCR analyses

For semi-quantitative RT–PCR, total RNA was extracted from one-week-old seedlings using TRIzol Reagent (ThermoFisher Scientific), following the manufacturer’s instructions. After treatment of two µg RNA with RQ1 RNase-free DNase (Promega) and reverse transcription using M-MLV Reverse Transcriptase (Promega) according to the manufacturer’s instructions, 16 ng was used for each PCR amplification. Primer pairs used for PCR and their sequences are listed in Supplemental Table S1.

### Mitomycin C sensitivity assay

For the Mitomycin C sensitivity assay, seeds were surface-sterilized and sown onto solid medium containing half-strength Murashige and Skoog salts, 1% sucrose, 0.8% agar and supplemented or not with 20 µM Mitomycin C (SIGMA). After stratification for 2 days at 4°C, plants were grown for two weeks and sensitivity analyzed by counting the number of true leaves as previously described [39, 103]. Plants with three true leaves or less were considered as sensitive [39, 103].

### Histochemical GUS staining for somatic homologous recombination assay

The well-characterized IU.GUS-8 and DGU-US-1 *in planta* recombination tester lines consisting of an interrupted *ß-glucuronidase* (*GUS*) gene were used to determine the frequency of somatic homologous recombination [68, 69].

For the SDSA/GC assay, transgenic plants carrying the transgene were crossed with the recombination tester IU.GUS-8 line [68]. Plants homozygous for the IU.GUS substrate loci and either wild-type or homozygous for the transgene were identified in the F2. Seeds from the F2 were used for recombination tests. For the SSA assay, wild-type lines homozygous for DGU.US substrate were transformed with the *pRAD51::DMC1g* expression cassette. T2 plants homozygous for both the DGU.US substrate and the *pRAD51::DMC1g* transgene were selected and used for the GUS assay. Wild-type control plants were also selected from the same transformation batch.

For histochemical GUS staining, seeds were surface-sterilized, stratified at 4°C for 2 days and grown in petri dishes on solid medium for 2 weeks. Seedlings were then harvested and incubated in staining buffer (0.2% Triton X-100, 50 mM sodium phosphate buffer (pH 7.2), 2 mM X-Gluc (5-bromo-4-chloro-3-indolyl-ß-D-glucuronic acid; Biosynth), dissolved in N,N-dimethylformamide). Plants were infiltrated under vacuum for 15 min and incubated at 37°C for 24 hours. Staining solution was then replaced with 70% ethanol to remove leaf pigments and blue spots were counted under a binocular microscope.

For MMC induction, 11-day-old seedlings were transferred on liquid medium supplemented with 20 µM MMC and further grown for 3 days before GUS staining.

### Meiotic chromosome spreads

Meiotic chromosome spreads were prepared according to [104]. Whole inflorescences were fixed in ice-cold ethanol/glacial acetic acid (3:1) and stored at −20°C until further use. Immature flower buds of appropriate size were selected under a binocular microscope and incubated for 75-90 min on a slide in 100µl of enzyme mixture (0.3% w/v cellulase (Sigma), 0.3% w/v pectolyase (Sigma) and 0.3% cytohelicase (Sigma)) in a moist chamber at 37°C. Each bud was then softened for 1 minute in 20 µl 60% acetic acid on a microscope slide at 45°C, fixed with ice-cold ethanol/glacial acetic acid (3:1) and air dried. Slide were mounted in Vectashield mounting medium with DAPI (1.5 µg.ml^-1^; Vector Laboratories Inc.).

### Immunocytological analyses

Slide preparation and immunostaining in root tip nuclei were performed as previously described [105], except that slides were incubated with RAD51 antibody (diluted 1:500) for 24 hours at 4°C and 48 hours in case of RAD51 / DMC1 (1:500; 1:20) double immunostaining. Immunolocalisation of proteins in pollen mother cells has been performed as described previously [106]. The anti-ASY1 raised in Guinea Pig was used at a dilution of 1:250 [107], the anti-ZYP1 raised in rabbit [108] was used at a dilution of 1:500, the anti-RAD51 raised in rat [19] was used at a dilution of 1:500, and the anti-DMC1 raised in rabbit [109] was used at a dilution of 1:20.

### Microscopy

All observations were made with a motorized Zeiss AxioImager.Z1 epifluorescence microscope using a PL Apochromat 100X/1.40 oil objective. Photographs were taken with an AxioCam Mrm camera and appropriate Zeiss filter sets. Image Z-stacks were captured in three dimensions (x, y, z) and further deconvoluted with the deconvolution module (theoretical PSF, iterative algorithm) of the Zeiss ZEN lite software. For presentation, pictures are collapsed Z-stack projections obtained using the Extended-focus module of the ZEN lite software.

The pixel intensity profile was analyzed via the colocalization module of the ZEN software using uncollapsed Z-stack files. The pixel intensity through a region of interest was extracted using the ZEN software and plotted against the X dimension. The quantitative co-localization analysis and calculation of Pearson’s correlation coefficient was performed using ImageJ modules Coloc2.

### Western blot analyses

Seven-day-old seedlings grown *in vitro* on 0.8% agar plates were transferred to liquid medium (1% sucrose and half-strength Murashige and Skoog salts medium) supplemented or not with 30 µM MMC and incubated for 8 hours. For total protein extraction, about 30 seedlings were frozen in liquid nitrogen and ground in 100 µL extraction buffer (0.1 M Tris-HCl pH 7.5, 20% Glycerol, 2 mM EDTA, 1 mM DTT, 0.2% NP40 Nonidet, 1X Complete Mini Protease inhibitor cocktail (Roche)) and centrifuged at 4°C for 30 min at 20 000 g. Total protein content in the supernatant was carefully determined using the Bradford assay (Bio-Rad). 30 µg of proteins were denatured in the presence of 4X Laemmli Buffer at 95°C for 5 min [110]. Proteins were electrophoresed on 10% SDS-polyacrylamide gels [110] and transferred to nitrocellulose membrane. Equal protein loading was carefully controlled by reversible ponceau staining of the nitrocellulose membrane. DMC1 proteins were detected with a rabbit DMC1 antibody at a dilution of 1:800 (diluted antibody in 3% w/v nonfat dry milk, 1XPBS, 0.1% Tween-20; overnight incubation at 4°C) followed by goat anti-rabbit HRP-conjugated secondary antibody (1:5000 dilution in 3% w/v nonfat dry milk, 1XPBS, 0.1% Tween-20 ; 1h incubation at room temperature) and electrochemiluminescence detection (Clarity Max ECL; Bio-Rad). Membrane was eventually imaged using ChemiDoc MP imaging system (Bio-Rad).

## Acknowledgments

We thank Mathilde Grelon, Peter Schlögelhofer and Chris Franklin for providing the DMC1 and ZYP1, RAD51, and ASY1 antibodies, respectively. We thank members of the recombination group for their help and discussions.

## Supporting information

**Supplemental Figure 1. DAPI staining of pollen mother cells (PMC) during meiosis in WT, *dmc1* and *dmc1* plants expressing the *pRAD51::DMC1g* transgene.**

In WT meiosis, full pairing and synapsis is observed during Pachytene. Chromosomes condense and five bivalents linked by chiasmata are visualized during Metaphase I. Homologous chromosomes segregate in Anaphase I and Meiosis II proceeds leading to the formation of four balanced nuclei. In contrast, *dmc1* mutants exhibit defects in pairing and synapsis in pachytene, and 10 univalents are visible at Metaphase I due to absence of chiasmata. Univalents segregate randomly in Anaphase I and this leads, after Meiosis II, to polyads. The meiotic defects of the *dmc1* mutant are complemented by the presence of *pRAD51::DMC1g*. (Scale Bar: 10 µm).

**Supplemental Figure 2. RT-PCR analysis of Homologous Recombination genes in seedlings after irradiation.**

RT-PCR expression analysis of *RAD51*, *DMC1*, *RAD54* and *HOP2/MND1* in 7-day-old untreated or gamma-irradiated seedlings expressing or not the *pRAD51::DMC1* transgene. Seedlings were irradiated at 100 Gy and expression was analyzed 2 hours after irradiation. Actin is used as a loading control.

**Supplemental Figure 3. Expression of *DMC1* in somatic cells has a dominant negative effect and does not substitute for RAD51 in DSB repair in somatic cells.**

**(A to E) Mitomycin C hypersensitivity of transgenic wild-type seedlings overexpressing *DMC1*.** Pictures of two-week-old seedlings grown without (**A**) or with 20 µM MMC **(B to D)** are shown. **(E**) Sensitivity of the seedlings was scored after 2 weeks and the fractions of sensitive plants (plants with less than 4 true leaves) are shown (3 biological repeats, each with N > 45 seedlings). Three independent WT lines expressing the *pRAD51::DMC1* transgene were tested (T1-1, T1-2 and T1-3).

**(F-G) MMC sensitivity of *RAD51-GFP* lines overexpressing *DMC1*. (F)** Pictures of two-week-old seedlings grown with 20 µM MMC. **(G)** Sensitivity of the seedlings was scored after 2 weeks and the fractions of sensitive plants (plants with less than 4 true leaves) are shown. Three independent *RAD51-GFP* lines overexpressing *DMC1* were tested (N > 45) and all three lines showed strong hypersensitivity to MMC. The RAD51-GFP fusion protein forms RAD51 filaments that support the activity of DMC1 in meiosis [39] but this is not sufficient for DMC1 mitotic activity.

**Supplemental Figure 4. Sequence alignment of DMC1 proteins** from *Saccharomyces cerevisiae* (ScDMC1) and *Arabidopsis thaliana* (AtDMC1) and the corresponding mutated AtDMC1-II3A variant. The alignment was generated using ClustalW. Numbers indicate amino acid positions. Red letters indicate the three conserved amino acids (R136A, R308A, K318A in Arabidopsis) that have been mutated in yeast to generated *dmc1-II3A*. Under the sequences, asterisks, colons and full stops indicate identical, conserved and semi-conserved residues respectively.

**Supplemental Figure 5. DMC1 protein expression in transgenic seedlings**

**(A)** DMC1g and DMC1-II3A protein are induced by MMC treatment. Total proteins were extracted from 1-week-old seedlings treated or not with 30 µM MMC for 8 hours and DMC1 abundance measured. No DMC1-specific band (37 kDa) is observed in untreated plants in both wild-type and transgenic plants. In contrast, while DMC1 is still absent in wild-type plants after DNA damage treatment, it becomes clearly visible in transgenic plants expressing either *DMC1g* or *DMC1-II3A*. **(B)** Ponceau Red staining of the above membrane before incubation with anti-DMC1 antibody, showing equal loading.

**Supplemental Figure 6. DMC1-II3A variant does not complement meiosis of *dmc1* mutant and does not interfere with RAD51-mediated repair of meiotic DBS.**

**(A)** Schematic representation of the *pDMC1::DMC1-II3A* construct. Exons are shown as blue rectangles. **(B)**Expression of the *pDMC1::DMC1II3A* in *dmc1* mutants do not restore fertility. Left panel : Wild-type plants have long siliques full of seeds, while *dmc1* mutants have very low fertility. Right panel shows the number of seeds per silique in DMC1, *dmc1* mutant and two *dmc1* + *pDMC1::DMC1-II3A* independent transformants. Mean is shown as black horizontal bar; N = 20 to 25 siliques per plant.

**(C)** DAPI staining of chromosomes during meiosis. Wild-type cells show pairing and synapsis of homologous chromosomes in pachytene, five bivalents at metaphase I and two groups of five chromosomes at anaphase I. *dmc1* mutants exhibit defective synapsis but ten intact univalent at metaphase I, which randomly segregate at anaphase I. A similar phenotype is observed in *dmc1* plants expressing *pDMC1::DMC1-II3A.* (Scale Bar: 10 µm).

**Supplemental Figure 7. Absence of DMC1 focus in transgenic plants expressing DMC1-II3A and impaired synapsis**

**(A)** Male meiocytes stained with ASY1 antibody (red) and DMC1 antibody (green). In wild-type meiocyte, numerous DMC1 foci are visible on chromosome axes. In contrast, *dmc1* plants expressing DMC1-II3A lack DMC1 focus. (Scale bar: 5µm). **(B)** Male meiocytes stained with the ASY1 antibody (red) and the ZYP1 antibody (green). In wild-type pachytene, ZYP1 extends along the entire length of the chromosomes. In *dmc1* plants expressing DMC1-II3A, ZYP1 staining is restricted to a few foci and short stretches. (Scale bar: 5µm).

**Supplemental Figure 8. Expression of DMC1-GFP fusion protein has a dominant negative effect and impairs RAD51-mediated repair of meiotic DBS.**

**(A)** Schematic representation of the *pDMC1::DMC1-GFP* construct. **(B)** Wild-type plants have long siliques, full of seeds, while *dmc1* mutants are nearly sterile. Expression of *pDMC1::DMC1-GFP* in *dmc1* mutants does not restore fertility and even strongly reduces fertility when expressed in *rad51+/−* or WT plants. **(C)** DAPI staining of chromosomes during meiosis. Wild-type cells show pairing and synapsis of homologous chromosomes in pachytene, five bivalents at metaphase I and two groups of five chromosomes at anaphase I. *rad51* mutants exhibit defective synapsis and extensive chromosome fragmentation. Meiotic spreads in DMC1+/+ RAD51+/+ plants expressing *DMC1-GFP* revealed a *dmc1-like* phenotype with defective synapsis but ten intact univalents at metaphase I, which randomly segregate at anaphase I. In contrast, extensive fragmentation is observed in DMC1+/+ RAD51+/− plants expressing *DMC1-GFP*. (Scale Bar: 10 µm).

**Supplemental Figure 9. Expression of DMC1-GFP fusion protein in meiocytes**

**(A) Immunolocalisation of DMC1 in *DMC1-GFP* plants.** Male meiocytes stained with DAPI (blue), anti-AtASY1 antibody (red), and anti-DMC1 antibody (green). Numerous DMC1 foci are visible on the chromosomes in WT plants expressing *DMC1-GFP* but not in *dmc1−/− _*DMC1-GFP. (Scale bar = 10 μm).

**(B-C) *In vivo* observation of DMC1-GFP fluorescence in meiocytes.**

DMC1-GFP fluorescence is observed in nuclei of DMC1-GFP transgenic plants. (Scale bar = 50 μm in B and 10 µM for close-up view in C).

**S1 Table. List of primers used in this study**

**S1 Data. Raw data for fertility, recombination test and RAD51 foci countings.** These are numerical data used for figures and that support the findings of this study.

## Notes

### Competing Interest Statement

The authors have declared no competing interest.

### Summary of Updates

Figures were updated.

